# Foraging guild modulates insectivorous bat responses to habitat loss and insular fragmentation in peninsular Malaysia

**DOI:** 10.1101/2023.01.03.522569

**Authors:** Quentin C.K. Hazard, Jérémy S.P. Froidevaux, Natalie Yoh, Jonathan Moore, Juliana Senawi, Luke Gibson, Ana Filipa Palmeirim

## Abstract

Despite mounting evidence on the ecological impacts of damming for biodiversity, little is known regarding its consequences in the hyper-diverse Southeast Asian tropical forests. Here we assess the effects of habitat loss and fragmentation on the diversity and activity of insectivorous bats within the hydroelectric Kenyir Lake in peninsular Malaysia. We surveyed bat assemblages on 26 islands and two mainland continuous forest sites using passive acoustic monitoring. Echolocation calls were classified into sonotypes, each corresponding to either one or multiple species, and grouped into foraging guilds. We then examined bat overall assemblage (sonotype richness, activity, and composition), guild- and sonotype-specific activity. From 9360 hours of recordings, we identified 16 bat sonotypes, including 10 forest (2854 bat passes), three edge (13 703) and three open-space foragers (3651). Sonotype richness increased towards denser forest structures (higher Normalized Difference Vegetation Index - NDVI), while species composition varied across the gradient of forest area. Forest foragers were positively affected by NDVI and negatively affected by distance to the closest neighbour, whereas edge foragers’ activity increased in smaller islands. Of the six sonotypes analysed, the activity of one forest sonotype increased with forest area, while that of one edge sonotype decreased. Ensuring habitat quality within insular forest remnants, in addition to their functional connectivity, maximises bat diversity, including the persistence of forest foraging species. Future hydropower development should therefore avoid the creation of a myriad of small, isolated, and habitat-degraded islands further characterised by altered levels of bat diversity and guild-level activity.

**Highlights:**

- We assessed the diversity of insectivorous bats in dam-induced islands in Malaysia
- Species persistence was modulated by island size and habitat quality
- Forest foragers were negatively affected by island isolation and degradation
- Edge foragers benefited from fragmentation, increasing in activity on smaller islands
- By creating multiple small, isolated, degraded islands, damming erodes bat diversity

## 1. Introduction

Humanity currently faces a need to reconcile human population growth, increasing energy demands, and the decarbonization of that energy. In this context, hydropower is an increasingly appealing option, representing 73% of the renewable energy produced in the world (The World Bank, 2016). Yet, river damming is also a major driver of habitat loss and insular fragmentation across lowland forests (Gibson et al., 2017). By flooding the lowland areas, dam construction often creates insular forest fragments matching the previous hilltops that become isolated within an inhospitable aquatic matrix (Lees et al., 2016). Although recent efforts have been made to understand the ecological consequences of hydropower (Palmeirim et al., 2022; Terborgh et al., 2009), few studies have targeted the Southeast Asian forests (Jones et al., 2016). Such understanding is therefore considered a priority for biodiversity conservation in the region (Coleman et al., 2019).

Species diversity persisting in insular forest fragments is typically affected by fragment size and isolation, which limit species population size and colonisation rates respectively (McArthur & Wilson, 1967). Human activities and edge effects affect habitat quality within insular fragments, thereby influencing remaining species diversity (Benchimol & Peres, 2015a). Species responses to habitat loss and insular fragmentation may further vary between (Palmeirim et al., 2022) and within biological groups (Brändel et al., 2020; Meyer & Kalko, 2008), as influenced by particular species traits (Meyer et al., 2008; Palmeirim et al., 2021). For instance, the persistence of mid-sized mammal species on islands can be related to their swimming capacity (Benchimol & Peres, 2015b), while that of lizard species is dictated by their thermoregulation mode (Palmeirim et al., 2017). Understanding the drivers of species persistence in insular fragmented landscapes – considering both environmental and intrinsic species characteristics – allows more efficient management actions to be proposed, which is not trivial given the expansion of the hydropower sector across hyper-diverse tropical forests (Couto & Olden, 2018).

Although habitat loss and insular fragmentation have been reported as important drivers of bat species’ local extinction in the Neotropics (Colombo et al., 2022; Meyer & Kalko, 2008), little is known for Asia. In fact, the only study targeting such effects on insectivorous bats in this region highlighted the importance of island area especially for forest-dependent species in East China (López-Bosch et al., 2021). In addition, in a non-insular matrix setting in peninsular Malaysia, the diversity of insectivorous bats was impacted by forest area, with species-specific responses being modulated by their habitat affinity (Struebig et al., 2008). Insectivorous bats emit echolocation calls to navigate their surroundings and locate food (Schnitzler et al., 2003). The characteristics of the calls produced, e.g., call shape, are adapted to a species’ foraging preferences (Denzinger & Schnitzler, 2013; Schnitzler & Kalko, 2001). For example, species adapted to foraging in the forest interior (forest foragers) use long constant frequency (CF) calls or very short, broadband, frequency modulated (FM) calls, adapted to particularly cluttered environments. Edge foragers use quasi-constant frequency (QCF) calls, or medium frequency calls (FMqCF) composed of an FM component followed by a short and quasi constant element (qCF), allowing them to locate and navigate between background features (medium frequency FM component), and to locate prey at an intermediate distance (qCF component). Open-space foragers use low frequency (LF) (<30 kHz) FMqCF calls with a narrow FM component and a long QCF component, enabling prey detection in vast empty spaces (Denzinger & Schnitzler, 2013; Schnitzler & Kalko, 2001). Bats of different foraging guilds are therefore expected to respond differently to habitat loss and fragmentation (Denzinger & Schnitzler, 2013; Schnitzler & Kalko, 2001). Within foraging guilds, bat calls are not species specific: the calls of several species have evolved in a convergent way to respond to analogous environmental pressures, resulting in very similar calls in species facing analogous ecological conditions and therefore preventing species-specific identification in certain instances (Gibb et al., 2019; Russo et al., 2017). To overcome this issue, bat calls are commonly classified into sonotypes, i.e., calls of similar shape and peak frequency (Roemer et al., 2021).

The study of insectivorous bats in Southeast Asia, especially forest and edge foragers, has been largely impaired by the ability of most insectivorous species to avoid live trapping methods, such as harp traps and mist nets (Kingston, 2013). With the increasing affordability of low-cost acoustic devices, insectivorous bat surveys are becoming more accessible and reliable, further allowing for high replication (Gibb et al., 2019; Hill et al., 2018). Utilising such technological advances, here we provide the first assessment of the effects of habitat loss and insular fragmentation on insectivorous bats in a dam-induced landscape in peninsular Malaysia. Using passive acoustic monitoring, we surveyed insectivorous bats in 26 forest islands and two mainland continuous forest sites. Across a gradient of habitat loss and insular fragmentation, we tested the effect of island size, isolation and habitat quality (island shape and NDVI), at the following levels of bat diversity (1) overall assemblage, considering sonotype richness, activity and assemblage composition, (2) foraging guild, separately considering the activity of forest, edge and open-space foragers, and (3) sonotype, given each sonotype activity. We predicted that small, isolated, and habitat-degraded insular forest patches would have a lower sonotype richness and lower activity, as well as a dominance of edge and open-space foragers. Conversely, we predicted greater species diversity and higher activity for forest sonotypes on large, high quality and less isolated forest patches and in the mainland continuous forest sites.

## 2. Material and methods

### 2.1 Study area

This study was conducted within the insular fragmented landscape of the Kenyir Lake and its surroundings in peninsular Malaysia. This artificial freshwater reservoir was formed in 1986 by the damming of the Kenyir river. The novel insular landscape occupies 260 000 ha and is composed of >340 islands ranging in size between 0.6 and 1428 ha embedded in the water matrix (Figure 1). Tropical humid forest on the islands and the adjacent mainland continuous forest are characterised by lowland and mid-elevation dipterocarp vegetation. The wider reservoir landscape was subject to selective logging prior to damming (Qie et al., 2011). This area experiences a wet season between November and March, and a dry season between May and October. Annual precipitation varies between 2700 and 400mm annually (Qie et al., 2011).

**Figure 1.**
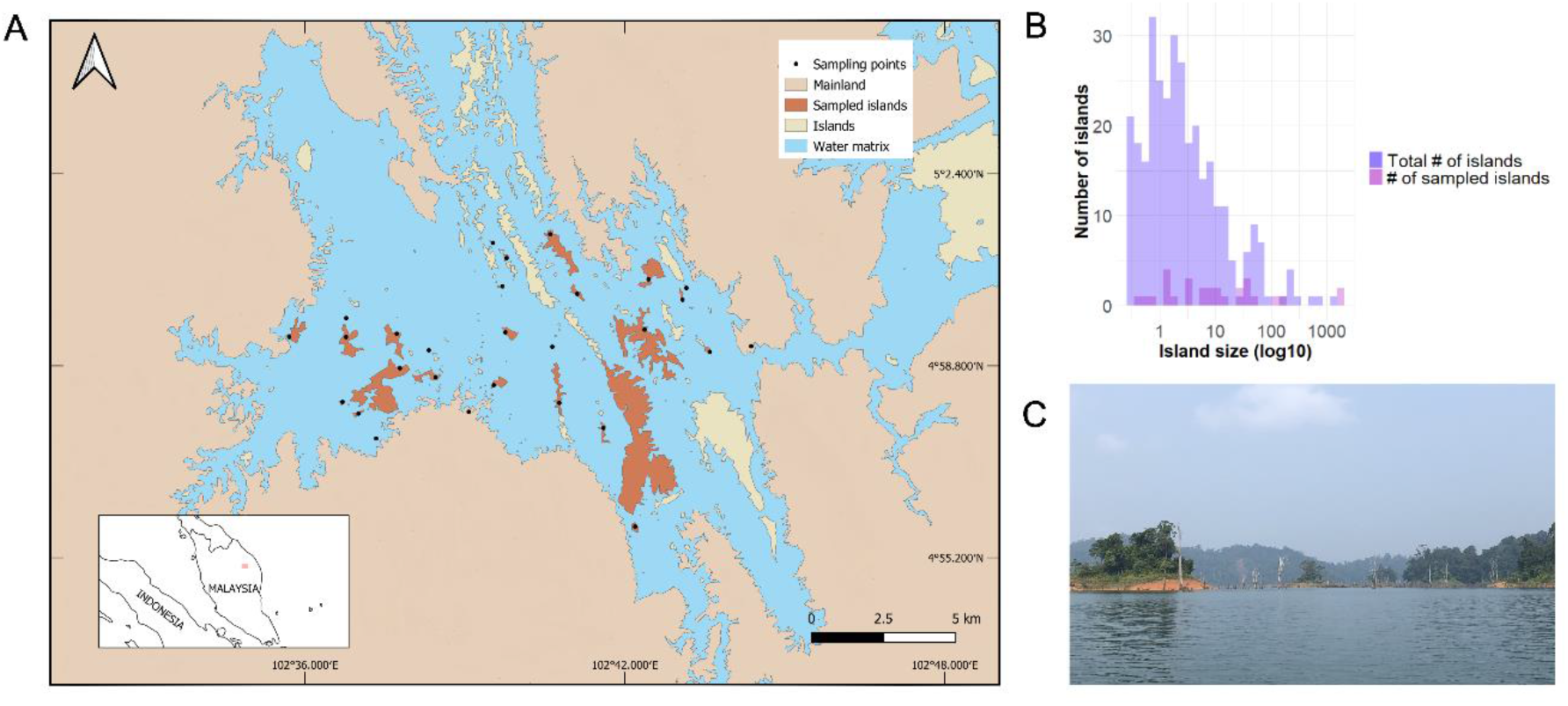
Map of the Kenyir Lake (A) Location of the study area and sampling sites in the Kenyir lake, peninsular Malaysia. The solid dots depict each of the 28 sampling sites. (B) Distribution of island sizes in Kenyir lake. (C) Photo of the Kenyir Lake.

### 2.2 Study design and data collection

We selected 26 islands covering a range of sizes (min-max: 0.45 – 167.3 ha) and degree of isolation from the mainland (135 – 2748 m), in addition to two mainland sampling sites. This sampling strategy was set up to study the effects of forest size and isolation independently, i.e. maintaining a low correlation between these variables (*r* = –0.36 when considering forest area and distance to mainland, *r* = – 0.49 when considering log-transformed forest area and distance to the mainland). Bat acoustic surveys were carried out between September 8^th^ and October 13^th^ 2019, using Audiomoth recorders (Hill et al., 2018) set on a sampling rate of 384 kHz and the gain to the second setting (“med”). We deployed one recorder in each sampling site and recorded for six hours. Recordings were divided into two time periods starting 30 minutes before sunset and ending 30 minutes after sunrise: from 18:00 to 22:00, and from 04:00 to 06:00 (Hayes, 1997), covering the two peaks of bat activity at dusk and dawn (Fenton, 1970). Each recorder was attached to a tree, positioned 2 metres above the ground and, to minimise any uncontrolled impact from edge effects, placed as inland as possible relative to island size, i.e., between 14 and 123 m from the edge (median: 50 m).

### 2.3 Acoustic analysis

Using the software Kaleidoscope Version 5.4.7 (Wildlife Acoustics, 2019), we split the recorded sequences into 5-second recordings (Torrent et al., 2018). The same software was used to filter the sequences containing sounds with a minimum frequency of 10 kHz and a maximum frequency of 250 kHz, and a pulse length between 2 and 500 ms. Among these sequences, only those containing one bat pass, i.e., at least two pulses of the same sonotype were kept for subsequent analysis (Torrent et al., 2018).

Prior to the acoustic analysis, we first compiled a list of all species of insectivorous bat known to occur in peninsular Malaysia (Lim et al., 2014; Nor Zalipah et al., 2019). Secondly, we collated reference calls for these species. We did so by conducting a literature survey using the Web of Science platform, between September and November 2021. We searched for publications by each species’ name followed by the country name: we favoured reference calls obtained in our study area in order to avoid any potential geographical variation in the call parameters. For those species we could not find any reference calls, we used the reference calls available in the bat call library Chirovox (Görföl et al., 2022). We therefore matched the call type of the species present in peninsular Malaysia to one of the sonotypes described in Yoh et al. (2022) for bat species in Malaysian Borneo, namely CF, FM, FMqCF1, FMqCF2, FMqCF3, FMqCF4, FMqCF5, QCF and LF sonotypes. Using start and end frequency, frequency of maximum energy, duration and interpulse interval as defined in Russo and Jones (2002), we classified the calls into one of these nine sonotypes. Given their very distinct echolocation parameters, the CF calls belonging to the genera *Rhinolophus* and *Hipposideros* could be identified to the species level.

As the shape of the echolocation calls reflects the physical constraints encountered by the bats, we were able to classify bat sonotypes into three foraging guilds: (1) the constant-frequency and FM calls represent forest foragers, (2) FMqCF4, FMqCF5 and QCF represent edge foragers, and (3) LF, FMqCF2 and FMqCF3 calls represent open-space foragers (Yoh et al., 2022). Social calls could not be identified to the sonotype level and were treated as assemblage-level activity.

### 2.4 Patch variables

Patch variables were obtained from a georeferenced LANDSAT 5 image which was transformed into a land/water matrix using an unsupervised classification on the software ArcGIS (ESRI, 2011). We then used the “landscapemetrics” R package (Hesselbarth et al., 2019) to extract: (1) island size (*area*; ha), (2) shortest Euclidean distance to the mainland (*dist*.*main*; m), (3) shortest Euclidean distance to the nearest neighbour island or mainland (*dist*.*neigh*; m), (4) island shape (*shape*), defined as the ratio between the patch perimeter and the hypothetical minimum perimeter of this patch, i.e. the perimeter of a maximally compact patch (McGarigal & Cushman, 2002), (5) the normalised difference vegetation index (*NDVI*), and (6) distance between the recorder and the forest edge (*dist*.*edge; m*). Given that the mainland continuous forest sites are characterised by an extensive forest coverage non-isolated area, we attributed these sites with the closest possible values to ‘reality’. This included area values of one order of magnitude higher than the largest island (1670 ha) and zero distances to either the mainland or the nearest neighbour. *NDVI* and *shape* were calculated as for the remaining sampling sites but considering a 1000 m buffer centred in the sampling site and excluding water. To streamline, we refer to the area of both islands and mainland size as ‘forest area’.

### 2.5 Data analysis

Assemblage-level metrics include sonotype richness, activity, and assemblage composition. Sonotype richness was defined as the number of sonotypes: this measure is representative of the diversity of call traits present at a site. Activity, i.e., the number of bat passes, was used as a proxy for bat abundance (Kunz et al., 2009). Assemblage composition was summarised as a single variable using a Non-Metric Multi-Dimensional Scaling (NMDS) ordination. This analysis was performed considering sonotype activity and using a Bray-Curtis similarity matrix (stress = 0.130). The scores of the first axis of the NMDS composed the assemblage composition metric. Guild activity was calculated by summing the activity of the individual sonotypes respectively belonging to the forest, open-space and edge guilds (Table S1). Sonotype-level responses were examined for the sonotypes recorded in more than 10 sites and which had more than 50 bat passes. This threshold was intended to ensure a normal distribution of the residuals, as well as homoscedasticity. Social and unidentified calls were only included in the assemblage-level analysis. Among the sonotypes that met the threshold to be analysed, namely FMqCF2, FMqCF3, FMqCF4, FMqCF5, LF, *R. trifoliatus, H. diadema* and QCF, *H. diadema* and FMqCF2 had unequal error variances and were therefore excluded from the analysis.

We first accounted for spatial autocorrelation by applying Mantel tests using the R package ‘ade4’ (Dray & Dufour, 2007). These tests correlate geographic distance between sampling sites and each response variable as well as the residuals of each model introduced in the subsequent section. We found no spatial autocorrelation (*p* > 0.05) in all instances. We also examined the pairwise correlation between patch variables using Pearson correlation coefficients. *Shape* and *area* (log_10_ *x*) (*r* = 0.720), as well as *area* (log_10_ *x*) and *NDVI* were highly correlated (*r* = 0.800). Given the overall importance of area explaining biodiversity patterns in insular forest fragments (Jones et al., 2021), we preferred to keep this metric to enable comparisons with other studies, whereas *shape* was excluded from subsequent analysis. Due to the lack of knowledge on the effects of canopy closeness on bats in this region, we chose to also keep *NDVI* in subsequent analyses. However, *area* (log_10_ *x*) and *NDVI* were not included together in a model. Collinearity between predictor variables was also examined using Variance Inflation Factors (VIFs), with no variable showing substantial collinearity (VIF>5) (Dormann et al., 2013).

We then analysed the combined effects of patch variables – *area, dist*.*main, dist*.*neigh* and *NDVI* – on (1) sonotype richness, activity and assemblage composition; (2) activity of forest, edge, and open-space sonotypes, and (3) the individual activity of eight sonotypes. To do so, we applied Linear Models (LMs) to each of these response variables, whose distribution was scrutinised prior to the analysis. The response variables regarding overall assemblage, guild and sonotype level activity, as well as forest area were log-transformed. Although all models were run with a gaussian error distribution, we initially considered a negative binomial distribution for the overall, guild-level and sonotype-level activity responses. Given that none of the models addressing individual sonotypes activity nor forest guild activity converged with a negative binomial error distribution, and that the distribution of these variables’ residuals was closer to a normal distribution when using a log-transformation with a gaussian error structure, we chose to retain that transformation and error structure in the models. We further considered *dist*.*edge* as a covariate in each model, aiming to control for any eventual effect of distance to the forest edge.

A candidate model set including all possible combinations of patch variables (including the covariate *dist*.*edge*), except combinations involving *area* (log_10_ *x*) and *NDVI* in the same model, was generated using the dredge function of the MuMIn R package (Barton, 2022). All models were ranked by Akaike Information Criteria corrected for small sample sizes (AICc: Burnham & Anderson (2002)). To account for model uncertainty in multi-model inference, we used a model-averaging approach considering the most parsimonious models, i.e. those having the lowest AICc within a ΔAICc <2 (ΔAICc = AICc_i_ − AICc_min_, *i* being the i^th^ model derived from the dredge) (Froidevaux et al., 2022). We report model average estimates along with their 95% confidence intervals (CIs) which were considered significant if not overlapping zero (Nakagawa & Cuthill, 2007). Assumptions about the normal distribution of the variables and their residuals were verified using the R packages “performance” (Lüdecke et al., 2021) and “Dharma” (Hartig, 2022). All data analyses were performed using R (Team R Development Core, 2020).

## 3. Results

In total, we recorded 21 197 bat passes from 16 different sonotypes: 10 forest, three edge and three open-space foragers (Table 1). Sonotype richness varied between 4 and 13 sonotypes per site, activity varied between 43 and 3351 bat passes per six hours recording. Activity varied greatly across sampling sites (43 – 3351, 757.03 ±744.18). While the edge forager FMqCF4 and the open-space forager LF were present at every site, the forest foragers *Rhinolophus refulgens, Hipposideros kunzi*, CF.46, *H. cervinus* and *H. bicolor* were only found at one location. *H. bicolor, H. cervinus* and *H. kunzi* were only found on the mainland, while CF.46, and *R. refulgens* were only found on one island each (Table S1). According to the NMDS, low values in the first axis were mostly associated with larger forest sites and forest foragers (*H. cervinus, H. kunzi, H. bicolor*, FM), while high values were associated with smaller islands, as well as with edge (FMqCF4, FMqCF5), open-space (FMqCF3), and forest foragers (*R. refulgens, R. affinis*) (Figure 2). Overall, 988 bat passes could not be identified to either the guild or to the sonotype level, including 981 bat passes corresponding to social calls (Table 1). *R. trifoliatus* was likely greatly influential on the response of the forest guild: being present in less than half the forest sites, it accounted for nearly 80% of the forest guild’s activity (Table S1).

**Table 1.**
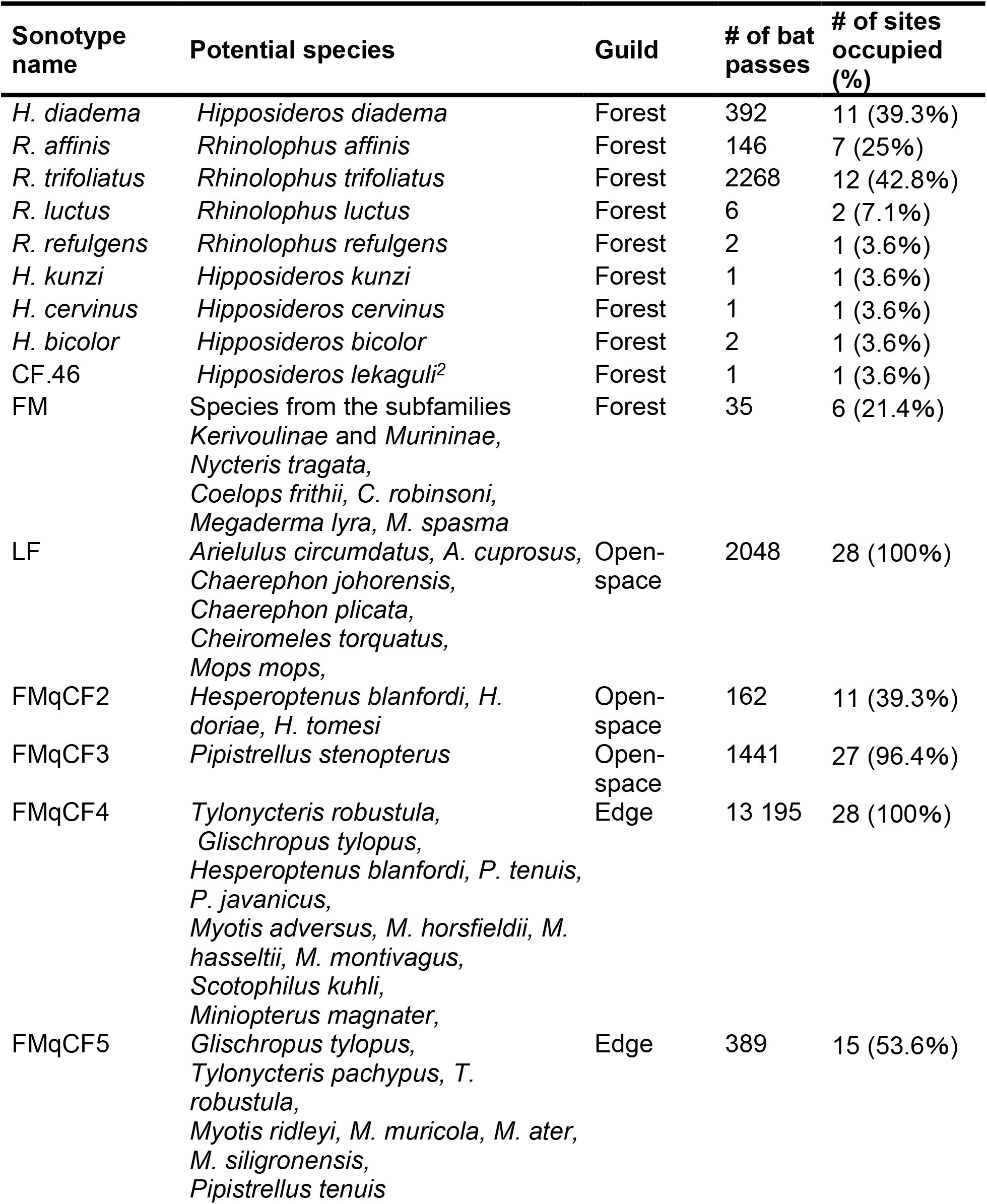
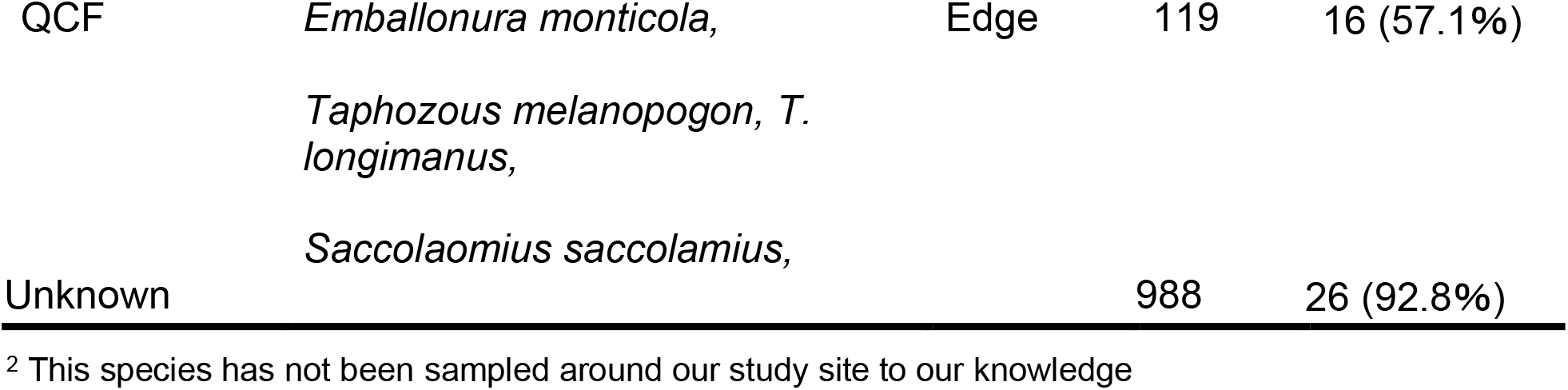
List of all sonotypes identified during the acoustic bat surveys across the insular fragmented landscape of the Kenyir Lake, peninsular Malaysia. For each sonotype, we indicate the potential species matching that sonotype, corresponding foraging guild, total activity (number of bat passes), and number of sampling sites in which each of the sonotypes was recorded. Bat passes that could not be identified were labelled as “unknown”.

**Figure 2.**
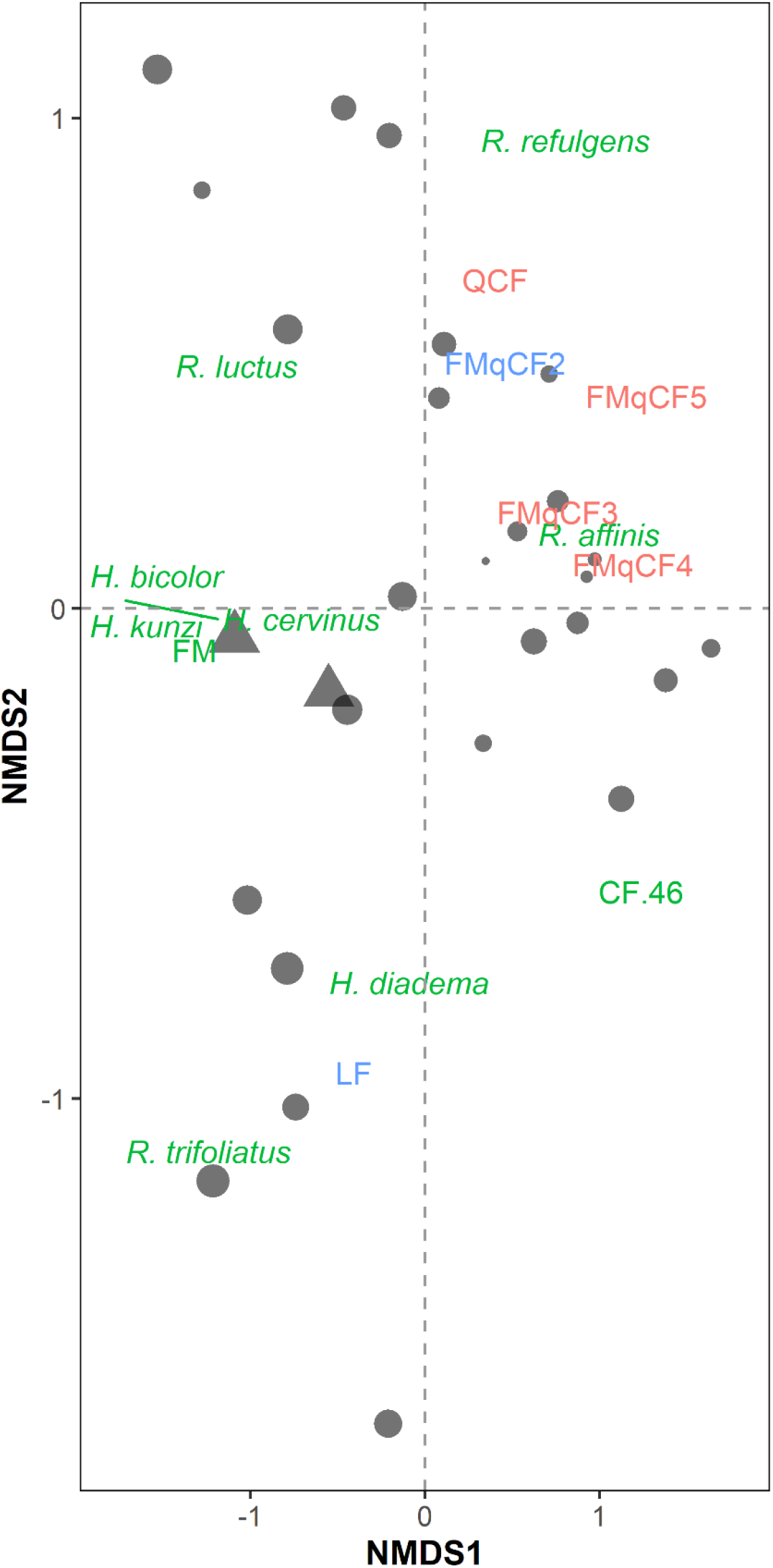
Non-Metric Multi-Dimensional Scaling (NMDS) ordination plot denoting both sampling sites and sonotypes. Sampling sites are represented by circles, matching the islands which are sized proportionally to their size (log_10_ *x*), and triangles correspond to the mainland continuous forest sites. Sonotypes are represented by their name and colour-coded according to the corresponding foraging guild: forest (in green), edge (red) and open-space (blue) (for further details regarding each sonotype, see Table 1). Given that the sonotypes *H. kunzi, H. cervinus* and *H. bicolor* are overlapping, for the sake of clarity, the position of these sonotypes is replaced by a green line and the sonotypes labels are separated.

### 3.1 Overall assemblage responses

Sonotype richness increased with NDVI (*β* = 0.819 ± 0.409, *p* = 0.045, CI _min_ = 0.017, CI _max_ = 1.620), while total bat activity was unaffected by the measured variables (Table S1). Assemblage composition varied among sites based on their size (*β* = – 0.536 ± 0.137, CI _min_ = –0.817, CI _max_ = –0.254), *p* < 0.001) (Table S1, Figures 2 and 3).

**Figure 3.**
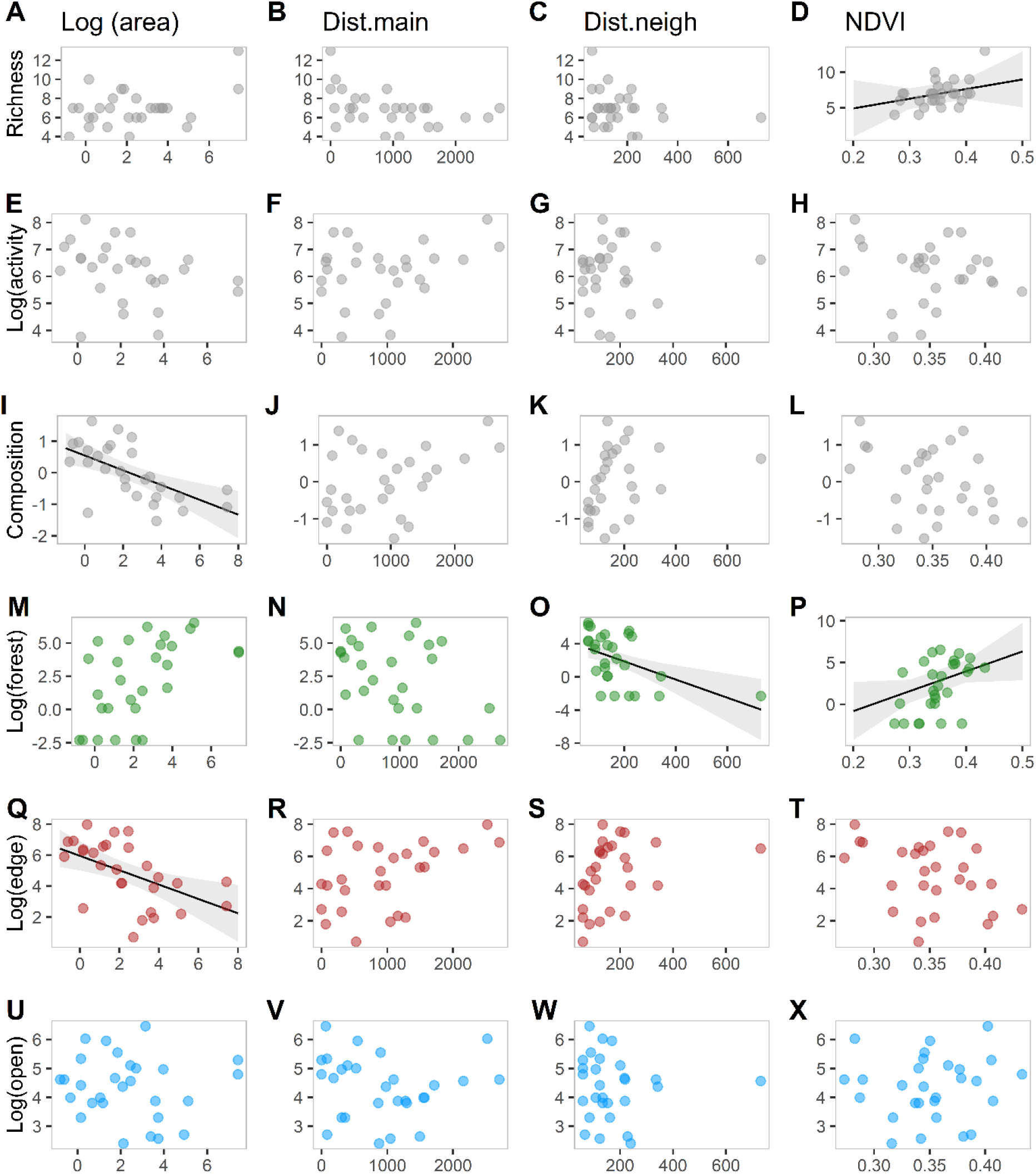
Relationships between bat sonotype richness (A-D), activity (log_10 *x*_) (E-H), assemblage composition (axis 1 of the NMDS) (I-L), and the activity of forest (log_10_ *x*) (M-P), edge (log_10_ *x*) (Q-T), and open-space foraging bats (log_10_ *x*) (U-X) and *Area* (log_10_ *x*) (A, E, I, M, Q, U), distance to the mainland (*Dist*.*main*) (B, F, J, N, R, V), Distance to neighbour (*Dist*.*neigh*) (C, G, K, O, S, W) and *NDVI* (D, H, L, P, T, X). In each panel, the solid black line is the prediction given by the averaged model obtained from the dredge, and the shaded area represents the 95% confidence interval. The predictions of the selected model were only shown for significant variables.

### 3.2 Guild-level responses

Forest sonotypes were more active at sites with higher NDVI (*β* = 1.476 ± 0.439, *p* = 0.002, CI _min_ = 0.616, CI _max_ = 2.335). The activity of forest sonotypes increased with decreasing distance to the closest neighbouring forest site (*β* = –1.468 ± 0.439, *p* = 0.002, CI _min_ = –2.328, CI _max_ = –0.608), while edge sonotypes activity decreased with increasing island size (*β* = –1.050 ± 0.366, *p* = 0.004, CI _min_ = –1.768, CI _max_ = – 0.332) and with distance to edge (*β* = –1.045 ± 0.355, *p* = 0.003, CI _min_ = –1.741, CI _max_ = –0.350). None of the tested variables had a significant effect on open-space sonotypes (Table S1, Figure 3). Unlike all other response variables, only one best model was selected for the activity of forest foragers (Table S2).

### 3.3 Sonotype-level responses

Among all the six individual sonotypes, only FMqCF4 and *R. trifoliatus* showed a significant response to the tested explanatory variables. FMqCF4 sonotype showed higher activity in smaller islands (*β* = –1.127 ± 0.365, *p* = 0.002, CI _min_ = –1.841, CI _max_ = –0.412), while *R. trifoliatus* were more active on larger forest sites (*β* = 1.487 ± 0.446, *p* = 0.001, CI _min_ = 0.614, CI _max_ = 2.361) (Table 2, Figure S1).

**Table 2.**
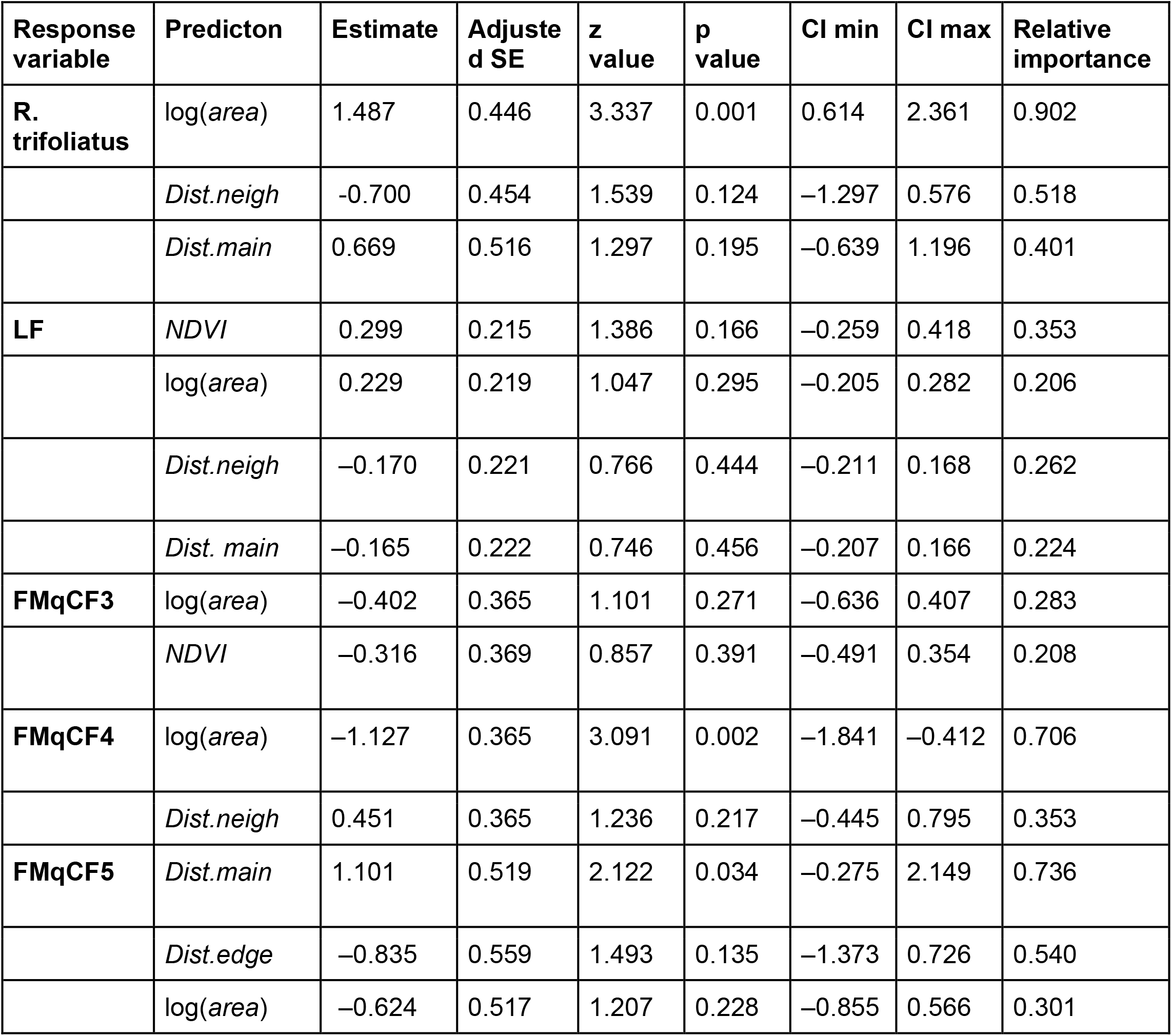
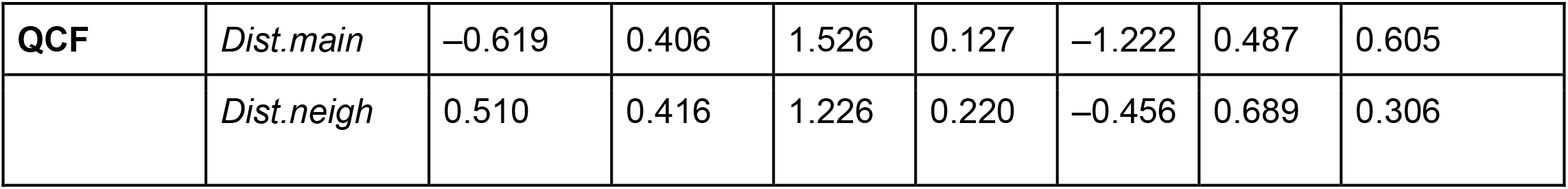
Standardized, model-averaged parameter estimates with associated standards errors (SE) and 95% confidence intervals (CIs) of the best LMs (ΔAICc < 2) relating the effects of forest area, isolation from the mainland (dist.main) and nearest neighbour (dist.neigh) and quality (NDVI) on bat activity at the sonotype level.

## 4. Discussion

A number of studies have demonstrated that habitat loss and insular fragmentation cause species local extinctions across lowland tropical forests (Gibson et al., 2013; Moore et al., 2022; Palmeirim et al., 2022; Pinto Henriques et al., 2021). Here, we contribute to fill an important knowledge gap by accordingly demonstrating overall negative bats response to disturbance across an insular fragmented landscape in Southeast Asia. Our results highlight the role of habitat quality driving the number of sonotypes, whereas forest area dictated which sonotypes were able to persist. Our guild-level analysis revealed that forest foragers were associated with greater habitat quality, particularly dense forest structures, and were negatively affected by increasing isolation from neighbouring landmasses. In contrast, edge foragers seemed to benefit from island shrinkage. Fragmentation effects were not so clearly observed at the sonotype-level, with only two of the six sonotypes analysed responding to patch variables, namely to forest area which had a positive effect on the forest forager *R. trifoliatus* and a negative effect on the edge sonotype FMqCF4.

Contrary to expectations given by the Island Biogeography Theory (McArthur and Wilson, 1967), forest area did not predict sonotype richness. Yet, forest area is an important driver of species diversity for several taxa in Amazonian reservoir islands (Jones et al., 2021), as well as for dung-beetles (Qie et al., 2011), primates and ungulates (Yong, 2015) surveyed in nearly the same islands in Kenyir. However, bat responses to patch size have been inconsistent across studies: in line with our results, no effect was observed for reservoir islands from the Brazilian Amazon and Panama (Colombo et al., 2022; Meyer & Kalko, 2008), but in China, López-Bosch et al. (2021) only observed a response in bat sonotype richness to island size after a certain size threshold was reached. However, Rocha et al. (2017) found that bat species richness increased at larger non-insular patches in the Brazilian Amazon. Despite the absence of clear effects of forest size, canopy closeness, as indicated by the NDVI, promoted an increase in the number of bat sonotypes across our study site. A bat species’ response to habitat quality is likely influenced by the intrinsic habitat characteristics such as 3D forest structure or canopy ruggedness, ultimately impacting which species are able to use each site (Froidevaux et al., 2016). The Kenyir Lake landscape has been subject to intensive selective logging prior to the construction of the dam (Qie et al., 2011). This activity that can lead to a loss of up to 80% of the canopy cover (Saiful & Latiff, 2019) resulted in an overall low, yet variable NDVI across the islands and surrounding continuous forest. Although the effects of logging on bat species richness seem to be limited both in the Neotropics (Meyer et al., 2016) and in the Paleotropics (Struebig et al., 2013), logging appears to strongly influence assemblage composition, edge species being indicative of repeatedly logged sites. Yet, the effects of logging on biodiversity depend on the intensity and extraction methods (Burivalova et al., 2014), and further investigations regarding the effects of logging intensity in the context of insular forest fragments are needed to further our understanding of how logging may drive bat sonotype richness.

In addition, forest insularisation led to the creation of edges, whose deleterious effects on vegetation include increased exposure to wind-throws, culminating in shifts towards disturbance-adapted pioneer trees (Benchimol & Peres, 2015a; Santo-Silva et al., 2021). While sites with low canopy closeness can be widely used by edge foragers, only those sites harbouring increased NDVI may represent suitable habitat for manoeuvrable forest dependent species that are further adapted to echolocate in more cluttered environments (Froidevaux et al., 2016; Suarez-Rubio et al., 2018). By allowing forest foragers to persist, habitat quality contributes to maintain bat diversity, as also observed for other biological groups, e.g., large-sized mammals and reptiles (Oliveira et al., 2020; Silva et al., 2022). This is further supported by the increase in forest bat activity we observed in denser forests. Higher NDVI values may also be associated with higher availability of mature trees that provide roosting sites for species such as *R. trifoliatus, R. sedulus, K. papillosa* and *K. pellucida*, all of which depend on these structures to rest and thus to persist. For instance, in Malaysia, the absence of tree cavities due to forest disturbance was associated with the decline of the forest foragers *Kerivoula* sp. (Struebig et al., 2013). Our findings reiterate the importance of habitat quality as a key driver of species diversity in fragmented landscapes (Armstrong et al., 2022; Poniatowski et al., 2018).

Furthermore, bat assemblage composition varied along the gradient of forest area, with edge foragers being particularly active on smaller islands. This trend was further reflected at the sonotype level by the edge forager FMqCF4. These responses were expected given that small islands tend to be edge-dominated. Furthermore, as a caveat to this study, in small islands, detectors had to be placed closer to edges given the lack of forest interior. It is therefore possible that the detectors on small islands, being mechanically closer to forest edges, recorded a higher activity of edge foragers. This is further supported by the negative relationship between edge foragers activity and the *Dist*.*edge*. In any case, this would still demonstrate the preferential use of edges by this bat guild (López-Bosch et al., 2021). In contrast, our results show that the forest forager *R. trifoliatus* responds positively to forest area, suggesting that this sonotype requires greater habitat complexity associated with larger areas of forest (Benchimol & Peres, 2015a).

Contrary to our expectations, isolation was not an important variable explaining bat assemblage-level responses. These results contrast with an insular fragmented landscape in Panama, where isolation to the mainland was the main predictor of bat richness (Meyer & Kalko, 2008). However, they support findings from a study in a non-insular Malaysian landscape, where isolation has also been found to be a poor prediction of bat richness (Brändel et al., 2020). This lack of isolation effect may be related to the overall small distance separating most of the study sites, and the overall size of the lake, as the home ranges for most local species exceed the distance separating most of the study sites (Wilson et al., 2010). Nevertheless, forest foragers were more active in sites less isolated from neighbouring landmasses, which might be due to morphological constraints (Norberg & Rayner, 1987). Indeed, forest foragers have a wing morphology characterised by a low aspect ratio (wingspan²/wing area) and a low wing loading (body mass/wing area) (Norberg & Rayner, 1987). Although this characteristic allows them to have a slow and highly manoeuvrable flight, it also makes flight over open spaces particularly energetically demanding (Altringham, 2011; Bader et al., 2015). Furthermore, the absence of distance-to-mainland effects in favour of distance-to-neighbour effects for forest foragers underlines the value of intermediary islands to act as stepping-stones for forest bats to cross the water matrix and reach more remote islands. This idea is supported by Saura et al. (2014) who also stress that these intermediate islands need to be sufficiently large and of high quality in order to act as stepping-stones.

Our results emphasise the valuable use of passive acoustic monitoring techniques to survey bat assemblages, further allowing us to examine bat responses at multiple levels. However, the use of sonotype richness instead of species richness likely biased the estimated sonotype richness towards forest species. Indeed, while the CF calls produced by forest foragers could be identified to the species level, other sonotypes including FMqCF, QCF and LF contained multiple species. Likewise, given the similarity of the ecological constraints faced by edge foragers, the calls produced by the species belonging to this guild can only be separated between three sonotypes (FMqCF4, FMqCF5 and QCF). For this reason, our results most likely underestimate the effects of habitat loss and insular fragmentation on insectivorous bats. In addition, as species detectability is a function of call intensity (Hayes, 2000), forest bats producing low-intensity FM calls such as Vespertilionidae (e.g., *Kerivoula* and *Myotis* spp.) tend to be under-detected (Waters and Jones, 1995). This might further explain the relatively weak responses observed at the sonotype-level, which should therefore be interpreted with caution. Live trapping remains the more efficient method to monitor these species (Kingston, 2013), and still, studies using these trapping methods have highlighted the high sensitivity of these forest genera to forest disturbance (Huang et al., 2019).

## 5. Conservation implications

Hydropower development is set to massively expand across Southeast Asian forests, with energy production expected to increase threefold by 2035 (Petinrin & Shaaban, 2015; Tang et al., 2019). In Malaysia alone, at least four additional major dams will soon be constructed (> 34 000 MW) (Foo, 2015). To minimise the detrimental effects of damming, our results suggest that special attention should be given to forest bats which, given their forest-adapted morphology, are particularly extinction prone (Jones et al., 2003; Safi & Kerth, 2004). In this context, our results emphasise that preserving forests with denser canopies that remain functionally connected to nearby landmasses is crucial to maintain forest species. Furthermore, the independent presence of these qualities does not guarantee the use by forest bats: only patches that are large, well connected to the mainland, and harbouring a high habitat quality can serve as stepping-stones, and therefore allow less vagile species to commute over the water matrix (Saura et al., 2014). Our findings should be considered in habitat management strategies aiming to minimise biodiversity loss across fragmented, lowland tropical forests. In insular forest patches, species are lost in a sustained and delayed manner according to the time elapsed since isolation, a process referred to as an “extinction debt” (Jones et al., 2016). Conservation efforts should prioritise maintaining habitat quality, for instance by minimising logging activity in highly forested areas (Hari Poudyal et al., 2018; Harvey & Brais, 2011). In addition, future hydropower developments should consider how dam placement is likely to affect the creation of different island systems. These developments should aim to reduce the creation of a myriad of small, isolated, and habitat-degraded forest which would help conserve biodiversity in these landscapes, for instance by targeting craggy locations, therefore minimising the flooded area.

## Abbreviations

AICc: Akaike Information Criterion corrected for sample size
CF: Constant Frequency
CI: Confidence Interval
FM: Frequency Modulated
FMqCF: Frequency Modulated quasi–Constant Frequency
LF: Low Frequency
LM: Linear Model
NDVI: Normalised Difference Vegetation Index
NMDS: Non-Metric Multi-Dimensional Scaling
QCF: Quasi-constant Frequency

