## Supplemental Table 1 for "Foraging guild modulates insectivorous bat responses to habitat loss and insular fragmentation in peninsular Malaysia"

**Supplementary material**

**Table S1.** Standardized, model-averaged parameter estimates with associated standards errors (SE) and 95% confidence intervals (CIs) of the best LMs (ΔAICc < 2) relating the effects of forest size, isolation and quality on bat response at the ensemble and guild levels.

| **Response variable** | **Predictor** | **Estimate** | **Adjusted SE** | **z value** | **p value** | **CI min** | **CI max** | **Relative importance** |
| --- | --- | --- | --- | --- | --- | --- | --- | --- |
| **Richness** | NDVI | 0.819 | 0.409 | 2.003 | 0.045* | 0.017 | 1.620 | 0.508 |
|  | Edge-point | 0.778 | 0.424 | 1.833 | 0.067 | –0.054 | 1.609 | 0.513 |
|  | Dist. to neighbour | –0.372 | 0.331 | 1.124 | 0.261 | –1.022 | 0.277 | 0.244 |
|  | Dist to mainland | –0.408 | 0.400 | 1.048 | 0.295 | –1.172 | 0.355 | 0.335 |
| **Activity** | log(area) | –0.309 | 0.213 | 1.453 | 0.146 | –0.726 | 0.108 | 0.291 |
|  | Dist to mainland | –0.297 | 0.213 | 1.392 | 0.164 | –0.121 | 0.715 | 0.344 |
|  | Edge point | –0.284 | 0.214 | 1.329 | 0.184 | –0.703 | 0.135 | 0.307 |
| **Assemblage composition** | log(area) | –0.536 | 0.144 | 3.729 | <0.001  *** | –0.817 | –0.254 | 0.895 |
|  | Dist to neighbour | 0.138 | 0.146 | 0.946 | 0.344 | –0.159 | 0.2437 | 0.286 |
| **Forest** | Dist to neighbour | –1.468 |  | –3.347 | 0.002  ** | –2.328 | –0.608 | 0.916 |
|  | NDVI | 1.476 |  | 3.364 | 0.002  *** | 0.616 | 2.335 | 0.675 |
| **Edge** | log(area) | –1.050 | 0.366 | 2.866 | 0.004** | –1.768 | –0.332 | 0.643 |
|  | Dist to neighbour | 0.411 | 0.355 | 1.158 | 0.247 | –0.285 | 1.108 | 0.324 |
|  | Dist.edge | –1.045 | 0.354 | 2.946 | 0.003** | –1.741 | –0.350 | 0.486 |
|  | Dist.main | 0.338 | 0.399 | 0.847 | 0.397 | –0.444 | 1.120 | 0.265 |
| **Open** | NDVI | 0.285 | 0.208 | 1.371 | 0.170 | –0.122 | 0.692 | 0.178 |
|  | Dist mainland | –0.224 | 0.211 | 1.061 | 0.289 | –0.637 | 0.189 | 0.224 |
