## Supplemental Table 2 for "Foraging guild modulates insectivorous bat responses to habitat loss and insular fragmentation in peninsular Malaysia"

**Table S2.** Best models (ΔAICc < 2) selected by the dredge procedure relating the effects of forest size, isolation and quality on bat response at the ensemble, guild and sonotype levels.

| **Response variable** | **Parameters** | **K** | **AICc** | **Delta AICc** | **AICc weight** |
| --- | --- | --- | --- | --- | --- |
| **Richness** | NDVI | 3 | 112.423 | 0.000 | 0.160 |
|  | Dist.edge | 3 | 12.509 | 0.085 | 0.154 |
|  | Dist. edge, NDVI | 4 | 113.508 | 1.084 | 0.093 |
|  | Dist.neigh, NDVI | 4 | 113.643 | 1.220 | 0.087 |
|  | NDVI, Dist.main | 4 | 113.855 | 1.432 | 0.078 |
|  | Dist.edge, Dist.main | 4 | 113.903 | 1.480 | 0.076 |
| **Activity** | log(area) | 3 | 87.301 | 0.125 | 0.137 |
|  | Dist.main | 3 | 87.491 | 0.315 | 0.125 |
|  | Dist.edge | 3 | 87.679 | 0.503 | 0.113 |
| **Assemblage composition** | log(area) | 3 | 64.357 | 0.000 | 0.390 |
|  | Dist.neigh, log(area) | 4 | 66.012 | 1.654 | 0.171 |
| **Forest** | Dist.neigh, NDVI | 4 | 132.129 | 0.000 | 0.404 |
| **Edge** | log(area) | 3 | 103.513 | 0.000 | 0.334 |
|  | Dist. neigh, log(area) | 4 | 104.998 | 1.485 | 0.159 |
| **Open** | NDVI | 3 | 85.994 | 0.379 | 0.156 |
|  | Dist. mainland | 3 | 86.833 | 1.218 | 0.103 |
| ***R. trifoliatus*** | log(area) | 3 | 123.394 | 0.000 | 0.251 |
|  | Dist.neigh, log(area), Dist.main | 5 | 123.837 | 0.444 | 0.201 |
|  | Dist. neigh, log(area) | 4 | 124.133 | 0.739 | 0.173 |
|  | log(area), Dist.main | 4 | 125.211 | 1.817 | 0.101 |
| ***H. diadema*** | Dist.edge | 3 | 120.731 | 0.000 | 0.265 |
|  | Dist.edge, NDVI | 4 | 121.412 | 0.681 | 0.188 |
|  | Dist.edge, log(area) | 4 | 122.393 | 1.661 | 0.115 |
| **FMqCF3** | log(area) | 3 | 117.582 | 1.119 | 0.130 |
|  | NDVI | 3 | 118.130 | 1.662 | 0.100 |
| **FMqCF4** | log(area) | 3 | 116.293 | 0.000 | 0.260 |
|  | Dist.neigh, log(area) | 4 | 117.205 | 0.912 | 0.165 |
| **FMqCF5** | Dist.main | 3 | 129.910 | 0.000 | 1.198 |
|  | Dist.edge, Dist.main | 4 | 130.832 | 0.931 | 0.124 |
|  | Log(area), Dist.main | 4 | 130.904 | 0.994 | 0.120 |
|  | Dist.edge | 3 | 131.794 | 1.882 | 0.077 |
| **QCF** | Dist.main | 3 | 118.092 | 0.217 | 0.154 |
|  | Dist.neigh, Dist.main | 4 | 119.032 | 1.157 | 0.097 |
| **LF** | NDVI | 3 | 88.062 | 0.331 | 0.152 |
|  | log(area) | 3 | 88.981 | 1.250 | 0.095 |
|  | Dist.neigh | 3 | 89.564 | 1.833 | 0.071 |
|  | Dist.main | 3 | 89.599 | 1.869 | 0.070 |
