## Supplementary figures and images for "Foraging guild modulates insectivorous bat responses to habitat loss and insular fragmentation in peninsular Malaysia"

### Supplemental Figure 1

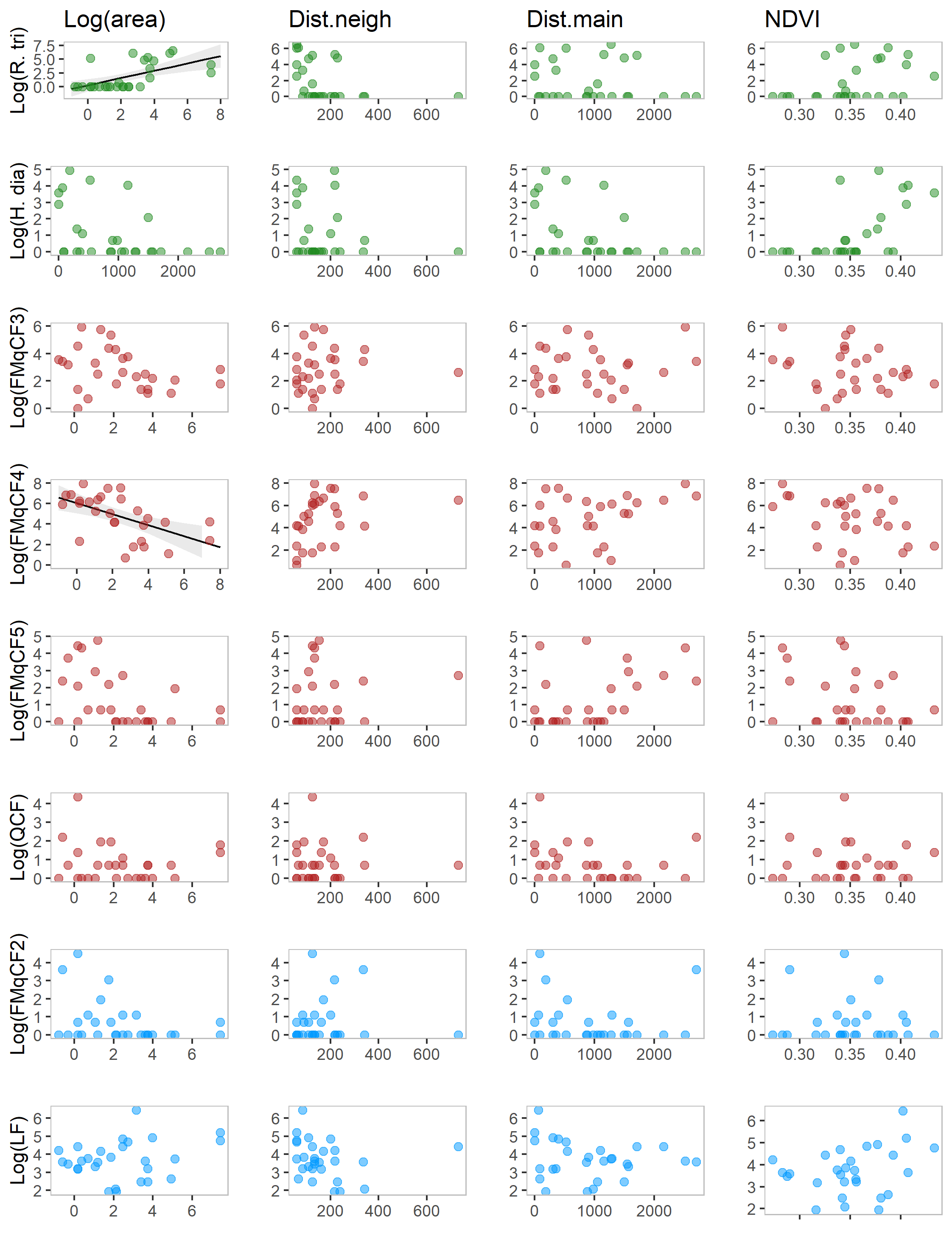
